# Mapping the subcortical connectivity of the human default mode network

**DOI:** 10.1101/2021.07.13.452265

**Authors:** Jian Li, William H. Curley, Bastien Guerin, Darin D. Dougherty, Adrian V. Dalca, Bruce Fischl, Andreas Horn, Brian L. Edlow

**Affiliations:** Center for Neurotechnology and Neurorecovery, Massachusetts General Hospital and Harvard Medical School, Boston, MA, USA; Athinoula A. Martinos Center for Biomedical Imaging, Massachusetts General Hospital and Harvard Medical School, Charlestown, MA, USA; Harvard Medical School, Boston, MA, USA; Department of Psychiatry, Massachusetts General Hospital and Harvard Medical School, Boston, MA, USA; Computer Science and Artificial Intelligence Laboratory, Massachusetts Institute of Technology, Cambridge, MA, USA; Department of Neurology, Movement Disorders and Neuromodulation Sectio Charite, University Medicine Berlin, Berlin, Germany

## Abstract

The default mode network (DMN) mediates self-awareness and introspection, core components of human consciousness. Therapies to restore consciousness in patients with severe brain injuries have historically targeted subcortical sites in the brainstem, thalamus, hypothalamus, basal forebrain, and basal ganglia, with the goal of reactivating cortical DMN nodes. However, the subcortical connectivity of the DMN has not been fully mapped and optimal subcortical targets for therapeutic neuromodulation of consciousness have not been identified. In this work, we created a comprehensive map of DMN subcortical connectivity by combining high-resolution functional and structural datasets with advanced signal processing methods. We analyzed 7 Tesla resting-state functional MRI (rs-fMRI) data from 168 healthy volunteers acquired in the Human Connectome Project. The rs-fMRI blood-oxygen-level-dependent (BOLD) data were temporally synchronized across subjects using the BrainSync algorithm. Cortical and subcortical DMN nodes were jointly analyzed and identified at the group level by applying a novel Nadam-Accelerated SCAlable and Robust (NASCAR) tensor decomposition method to the synchronized dataset. The subcortical connectivity map was then overlaid on a 7 Tesla 100 micron *ex vivo* MRI dataset for neuroanatomic analysis using automated segmentation of nuclei within the brainstem, thalamus, hypothalamus, basal forebrain, and basal ganglia. We further compared the NASCAR subcortical connectivity map with its counterpart generated from canonical seed-based correlation analyses. The NASCAR method revealed that BOLD signal in the central lateral nucleus of the thalamus and ventral tegmental area of the midbrain is strongly correlated with that of the DMN. In an exploratory analysis, additional subcortical sites in the median and dorsal raphe, lateral hypothalamus, and caudate nuclei were correlated with the cortical DMN. We also found that the putamen and globus pallidus are negatively correlated (i.e., anti-correlated) with the DMN, providing rs-fMRI evidence for the mesocircuit hypothesis of human consciousness, whereby a striatopallidal feedback system modulates anterior forebrain function via disinhibition of the central thalamus. Seed-based analyses yielded similar subcortical DMN connectivity, but the NASCAR result showed stronger contrast and better spatial alignment with dopamine immunostaining data. The DMN subcortical connectivity map identified here advances understanding of the subcortical regions that contribute to human consciousness and can be used to inform the selection of therapeutic targets in clinical trials for patients with disorders of consciousness.

## 1. Introduction

Recent advances in structural and functional connectivity mapping create opportunities for therapeutic neuromodulation of human brain networks (Horn and Fox, 2020). For patients with disorders of consciousness (DoC) caused by severe brain injuries, functional connectivity mapping can be used to identify widely connected network hubs that are therapeutic targets for stimulation (Edlow et al., 2020). The biological and mechanistic rationale for this targeted approach to neuromodulation has been demonstrated in rodent (Taylor et al., 2016) and non-human primate (Redinbaugh et al., 2020) models, which show that stimulation of subcortical network hubs promotes cortical reactivation and reemergence of consciousness from anesthetic coma. Emerging evidence from human electrical (Corazzol et al., 2017; Schiff et al., 2007; Thibaut et al., 2014), pharmacologic (Giacino et al., 2012; Whyte et al., 2014), and ultrasound-based (Cain et al., 2021a, 2021b; Monti et al., 2016) stimulation studies provide proof of principle that promoting recovery of consciousness in patients with DoC is possible (Edlow et al., 2021a). However, consensus on optimal subcortical therapeutic targets for neuromodulation of consciousness in humans has not been established (Edlow et al., 2021b).

A promising approach to identifying subcortical therapeutic targets is a “top-down” analysis of functional connectivity from canonical cortical networks that sustain consciousness. It is well established that the default mode network (DMN) contributes to self-awareness in the resting, conscious human brain (Buckner and DiNicola, 2019; Qin and Northoff, 2011; Raichle and Snyder, 2007). Although DMN functional connectivity alone is not sufficient for consciousness (Bodien et al., 2019; Demertzi et al., 2015; Norton et al., 2012), dynamic interactions between DMN nodes in the posterior cingulate, precuneus, medial prefrontal cortex, and inferior parietal lobules appear to be primary contributors to the neural correlates of consciousness (Bodien et al., 2017; Edlow et al., 2021a; Koch et al., 2016).

Given that the cortical nodes of the DMN are well-characterized and could be directly targeted with noninvasive methods such as transcranial direct current stimulation (tDCS) and transcranial magnetic stimulation (TMS), one could ask why the subcortical nodes of the DMN are clinically relevant. We could extend the question and ask why invasive stimulation methods such as deep brain stimulation (DBS) are not applicable at the cortical level (e.g., by placing a DBS electrode into the precuneus). Crucially, invasive methods have targeted subcortical regions for good reason, which has been referred to as the “funnel effect” of smaller brain nuclei (Parent and Hazrati, 1995). Projecting from cortical to subcortical structures (Swanson, 2000), information dimensionality (which is decompressed and openly available on cortical levels) is reduced (Bar-Gad et al., 2003). In ascending loops from subcortex to cortex, the reverse happens: information is expanded, decompressed, or de-referenced (Blouw et al., 2016). This architectural feature of the brain (Bota et al., 2015), which involves high-low-high dimensionality transforms of information, renders effects of neuromodulation on cortical versus subcortical levels strikingly different. Based on the large receptive and projective fields of subcortical brain structures, targeted neuromodulation of a small nucleus will affect a widely distributed and surprisingly large fraction of the entire cortex (Horn et al., 2019, 2017; Schiff et al., 2007). In contrast, diffuse neuromodulation techniques (e.g., TMS, tDCS), which modulate broad patches of cortical tissue, could have similar effects on networks (Fox et al., 2014). But doing the reverse (e.g., TMS to subcortical regions or DBS to cortical regions) would likely not produce the desired therapeutic effects. Hence, we believe it is crucial to precisely define subcortical nodes of the DMN to restore consciousness and cognitive function using targeted neuromodulation approaches such as DBS and low-intensity focused ultrasound pulsation (LIFUP).

Preliminary studies suggest that specific subcortical nuclei within the thalamus (Alves et al., 2019; Cunningham et al., 2017; Lee and Xue, 2018), basal forebrain (Alves et al., 2019), midbrain (Bär et al., 2016), pons (Fransson, 2005), and striatum (caudate and putamen) (Choi et al., 2012; Di Martino et al., 2008) are structurally and functionally connected to cortical DMN nodes. Testing for functional connectivity between subcortical regions and cortical DMN thus provides an opportunity to identify subcortical therapeutic targets in patients with DoC. Many such targets are amenable to therapeutic modulation by electromagnetic (Elias et al., 2020; Gratwicke et al., 2013; Kakusa et al., 2020; Schiff et al., 2007) and ultrasound-based therapies (Cain et al., 2021a, 2021b; Monti et al., 2016). However, DMN subcortical connectivity has not been fully mapped. In large part, this gap in knowledge is attributable to insufficient spatial resolution and low signal-to-noise ratio (SNR) of functional MRI, which poses a significant challenge to mapping functional connectivity for individual subcortical nuclei (Lee and Xue, 2018; Sclocco et al., 2018).

In this study, we aimed to create a comprehensive map of the subcortical connectivity of the DMN by combining high-resolution functional and structural datasets with advanced signal processing methods. Specifically, we used the resting-state functional MRI (rs-fMRI) dataset from 168 subjects acquired at 7 Tesla (7T) within the Human Connectome Project (HCP) (Smith et al., 2013). The rs-fMRI BOLD data were temporally synchronized across subjects using the BrainSync algorithm (Akrami et al., 2019; Joshi et al., 2018), which aligned all subjects’ data into the same spatiotemporal space, making it possible to model brain networks as low-rank components. The cortical and subcortical data were jointly analyzed and a more complete DMN (which spans both cortex and subcortex) was identified at the group level by applying a novel Nadam-Accelerated SCAlable and Robust (NASCAR) tensor decomposition method (Li et al., 2019b, 2021). The subcortical functional connectivity map was then overlaid on the 7T 100 micron *ex vivo* MRI dataset (Edlow et al., 2019) for precise neuroanatomic analyses of the brainstem, thalamus, hypothalamus, and basal ganglia using the FreeSurfer segmentation atlas (Fischl, 2012), probabilistic thalamic segmentation atlas (Iglesias et al., 2018), the Harvard ascending arousal network atlas (Edlow et al., 2012), and the basal forebrain and hypothalamus atlas proposed in (Neudorfer et al., 2020).

We first tested the hypothesis that the central lateral nucleus (CL) of the thalamus and the ventral tegmental area (VTA) of the midbrain are strongly connected to cortical DMN. This hypothesis is based on evidence from anatomic connectivity studies (Alves et al., 2019; Morales and Margolis, 2017; Schiff, 2010, 2008; Yetnikoff et al., 2014), animal neuromodulation studies (Baker et al., 2016; Redinbaugh et al., 2020; Solt et al., 2014; Taylor et al., 2016), and limited human studies (Schiff et al., 2007), which collectively indicate that CL and VTA are widely connected subcortical network hubs whose stimulation may activate the cerebral cortex and promote reemergence of consciousness. Second, we performed exploratory analyses to identify additional subcortical regions whose BOLD signal shows strong positive or negative correlation (i.e., anti-correlation) with the cortical DMN, indicating that these regions could potentially be used as alternative targets of neuromodulation. Finally, we explored the functional connectivity differences between the NASCAR approach and the traditional seed-based method and compared the results to immunostain data from a human brainstem specimen. We release the subcortical DMN functional connectivity map via the <u>Lead-DBS</u>, <u>FreeSurfer</u> and <u>Openneuro</u> platforms for use in future neuromodulation studies.

## 2. Materials and methods

### 2.1. In-vivo 7T resting-state fMRI data

We analyzed 7T resting-state fMRI (rs-fMRI) scans of healthy volunteers available from the Wash U/U Minn component of the Human Connectome Project (HCP) (Van Essen et al., 2012). We chose 7T, instead of 3T, dataset as it provides better SNR, particularly in subcortical regions. Eight subjects were excluded due to acquisition and/or preprocessing issues according to the HCP data release update (HCP Wiki, 2020), resulting in a total of 168 subjects used in this study. These 168 subjects were randomly split into two equally sized groups for reproducibility analysis. The following experiments were carried out on each group independently (84 subjects in each group). The rs-fMRI data were collected in four independent sessions with opposite phase encoding directions (PA, AP) using a gradient-echo EPI sequence (1.6 mm^3^ isotropic voxels, TR = 1000 ms, TE = 22.2 ms), where each session was 15 mins long (*T* = 900 frames). Only the first session (PA) was used in this work to minimize the potential inter-subject misalignment due to the different EPI distortions in different phase encoding directions, although EPI distortion had been carefully corrected during the preprocessing (Smith et al., 2013). The analyses were performed on the HCP minimally preprocessed 7T rs-fMRI data (Glasser et al., 2013), which were resampled and coregistered onto a common atlas in MNI space. The data were then represented in a grayordinate system (Glasser et al., 2013), where there are approximately 32K vertices on each hemisphere for cortical data and approximately 32K voxels for subcortical data. No additional spatial smoothing beyond the standard minimal preprocessing pipeline (2 mm full width half maximum (FWHM) isotropic Gaussian smoothing) was applied, because linear smoothing often blurs boundaries between different functional regions (Bhushan et al., 2016; Li et al., 2018, 2020a; Li and Leahy, 2017), which is problematic in resolving the relationships between small subcortical sub-regions in the brainstem, thalamus, hypothalamus, and basal forebrain.

### 2.2. Inter-subject temporal synchronization

Resting-state fMRI data are not directly comparable between subjects, as spontaneous BOLD activities in different subjects are not temporally synchronized. This is a critical issue even in stimulus-locked task fMRI data, where identical task design is used, because response latencies may differ between subjects (Friston et al., 1998). However, one of the assumptions in the low-rank tensor model we used in this work (described in the next section) is temporal synchrony across subjects, as the model does not work well on asynchronous fMRI data (Li et al., 2021). Therefore, we applied the BrainSync algorithm to achieve temporal alignment of the fMRI data (Joshi et al., 2018). BrainSync seeks an optimal temporal orthogonal transformation between two subjects, such that after synchronization the time series in homologous regions of the brain are highly correlated. In order to avoid the potential bias introduced by selecting any specific reference subject, we used the extended group BrainSync algorithm (Akrami et al., 2019) to build one virtual reference subject. This virtual reference subject is close, in the mean square sense, to all real subjects in the high dimensional space. Then we aligned all real subjects’ data to that virtual reference to obtain a multi-subject synchronized dataset, Fig. 1 (a). Crucially, applying BrainSync will not alter functional connectivity metrics (as carried out by correlation coefficients across BOLD series) when calculated using the whole time period (Joshi et al., 2018).

**Fig. 1.**
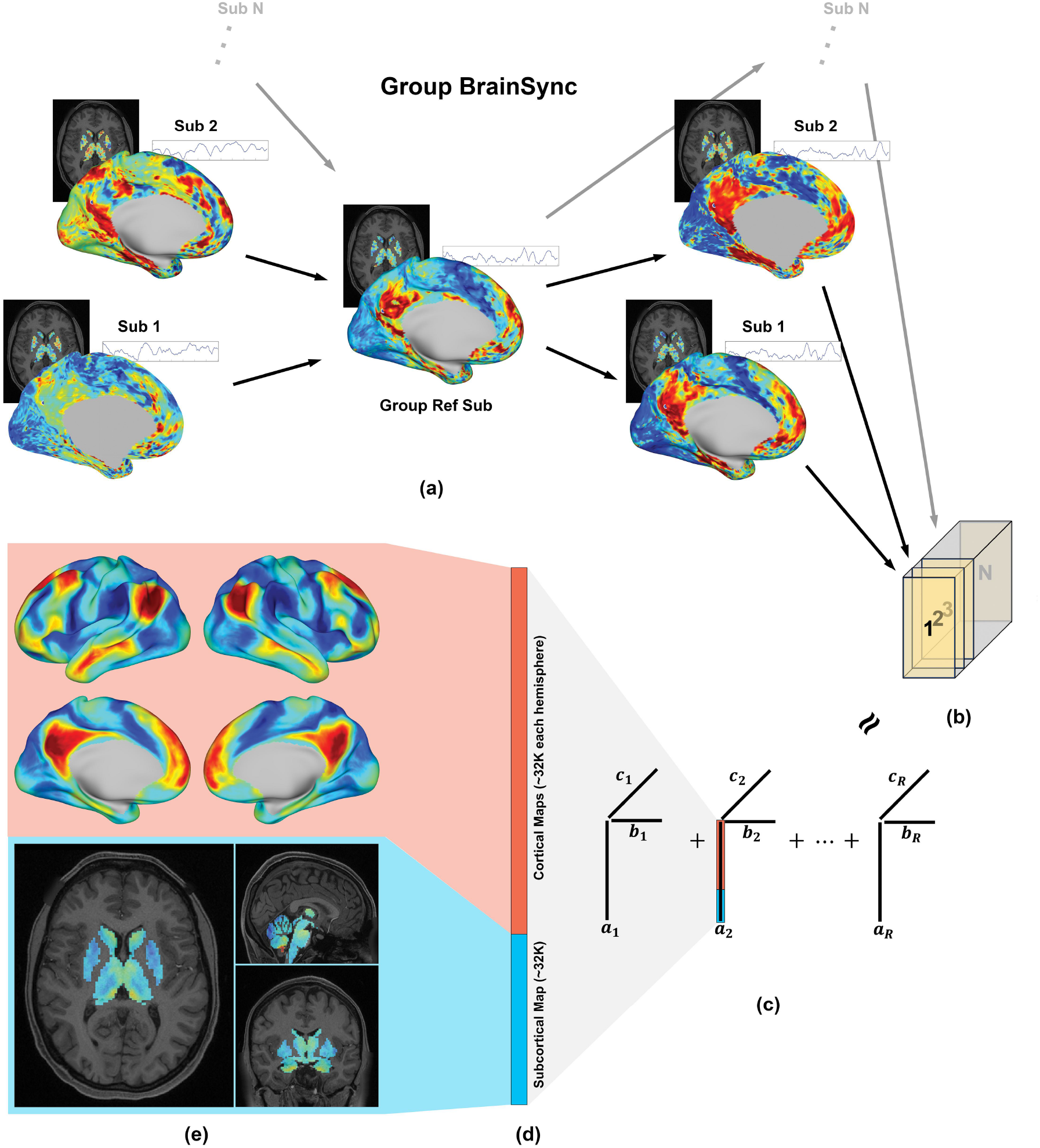
Brain network identification pipeline. (a) Group BrainSync transform for temporal alignment; (b) 3D tensor formation (space x time x subject); (c) Tensor decomposition using the Nadam-Accelerated SCAlable and Robust (NASCAR) canonical polyadic decomposition. ***a**_i_*, ***b**_i_*, and ***c**_i_* are the spatial map, the temporal dynamics, and the subject participation level for *i*^th^ component, respectively; (d) Grayordinate representation of the spatial map of the default mode network (DMN); (e) The cortical map and the subcortical map of the DMN.

### 2.3. Tensor-based brain network identification

Let 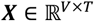 be the synchronized rs-fMRI data of an individual subject, where *V* is the number of vertices or voxels (space) and *T* = 900 is the number of time points (time). All subjects were concatenated along the third dimension (subject), forming a data tensor 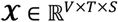, where *S* is the number of subjects, Fig. 1 (c). We model brain networks present in the group rs-fMRI data as a low-rank Canonical Polyadic (CP) model. Mathematically, the tensor 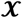 can be expressed as a sum of *R* rank-1 components:

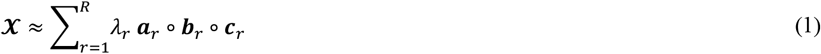

where each rank-1 component *λ_i_ **a**_i_* ∘ ***b**_i_* ∘ ***c**_i_* can be viewed as a brain network; 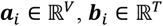 and 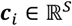 are the spatial map, the temporal dynamics, and the subject participation level, respectively, in the *i*^th^ network; *λ_i_* is the magnitude of that network, representing a relative strength of the activity in the *i*^th^ network to other networks; “∘” represents the outer product between vectors. *R* is the desired total number of networks. We use *R* = 30, as the rank of fMRI data has been shown to be limited (Calhoun et al., 2001; Li et al., 2021), and the network identification result is not sensitive to the choice of *R* as we explain below (Li et al., 2019b, 2021).

We solved this network identification (tensor decomposition) problem (Eq. (1)) using the Nadam-Accelerated SCAlable and Robust (NASCAR) canonical polyadic decomposition algorithm (Li et al., 2019b, 2021). NASCAR employs an iterative method using low-rank solutions as part of the initializations when solving higher-rank problems. The robustness of the solutions to initializations and the choice of *R* and the scalability to large dataset is substantially improved by using this warm start approach, and its superior performance over other traditional network identification methods has been demonstrated in applications to both electroencephalography (EEG) data (Li et al., 2019a, 2017) and fMRI data (Li et al., 2021, 2020b, 2019b).

### 2.4. Visualization of subcortical DMN connectivity

The DMN was identified as the second strongest network (second largest *λ* value), with the “physiological” signal being the strongest network (see the Discussion section). This is expected, as the DMN has been shown to be the most prominently active (or stable) brain network at rest (Buckner and DiNicola, 2019; Raichle, 2015). In contrast to the traditional definition of the DMN where only cortical nodes are considered, we redefine the DMN as a functional brain network that spans both cortex and subcortex. To explore the DMN subcortical functional connectivity, the subcortical section of the spatial DMN map was separated from the cortical section using grayordinate indices, Fig. 1 (c) and (d). The cortical section of the DMN, which we refer to as the “cortical map”, was plotted on the tessellated (inflated) surfaces for reference, as shown at the top of Fig. 1 (e). The subcortical counterpart was converted into a 3D volumetric representation in MNI space, referred to as the “subcortical map”, and trilinearly interpolated into a 0.5 mm^3^ isotropic resolution using the “mri_convert” tool in FreeSurfer (Fischl, 2012), as shown at the bottom of Fig. 1 (e). This interpolation procedure to higher resolution enabled both better visualization and the region-of-interest (ROI)-based analyses described below. Small ROIs in the atlases, such as CL, could vanish if the atlases are downsampled to the rs-fMRI resolution.

For better visualization of subcortical functional connectivity, particularly in relation to subcortical anatomy, the subcortical map was overlaid on a 7T 100 micron resolution *ex vivo* MRI dataset (Edlow et al., 2019) for precise neuroanatomic analyses. This dataset was acquired using a customized 31-channel coil over 100 hours of scan time and was co-registered to the MNI space. A manual examination and minor registration adjustment were performed to account for the subtle difference in the MNI template used for registering functional data in the HCP pipeline and that used in registering the 100 *μm* structural dataset.

### 2.5. Region-of-interest-based analysis

To study the functional connectivity for each subcortical ROI, we performed segmentation of the subcortical structures on the minimally preprocessed (Glasser et al., 2013) T1-weighted image of a reference subject (HCP subject ID: 100610, the default 7T subject provided by the HCP) using the automated segmentation tool (aseg atlas) in FreeSurfer (Fischl, 2012). Further sub-division of the thalamus and segmentation of thalamic nuclei was performed using a probabilistic atlas (PTN atlas) (Iglesias et al., 2018). The default segmentation output was used for all thalamic nuclei except for CL. Considering the small size and irregular shape of the CL nucleus, the CL segmentation mask was obtained by thresholding its posterior probability map at 0.03 (88^th^ percentile). This threshold was estimated by visual inspection of the averaged CL map over 100 HCP subjects so that the thresholded map accurately represents the shape and location of the CL nucleus. We used the Harvard ascending arousal network atlas (AAN atlas) (Edlow et al., 2012) for sub-division of brainstem nuclei, and an atlas proposed in (Neudorfer et al., 2020) for sub-division of basal forebrain and hypothalamus (BF/HT) nuclei. Labels of the AAN atlas were manually traced and provided in the MNI space (Edlow et al., 2012). Finally, all atlases were upsampled into a 0.5 mm^3^ isotropic resolution if the original atlases were in a lower resolution. Details about the ROIs with respect to the atlases are shown in Table 1.

**Table 1.**
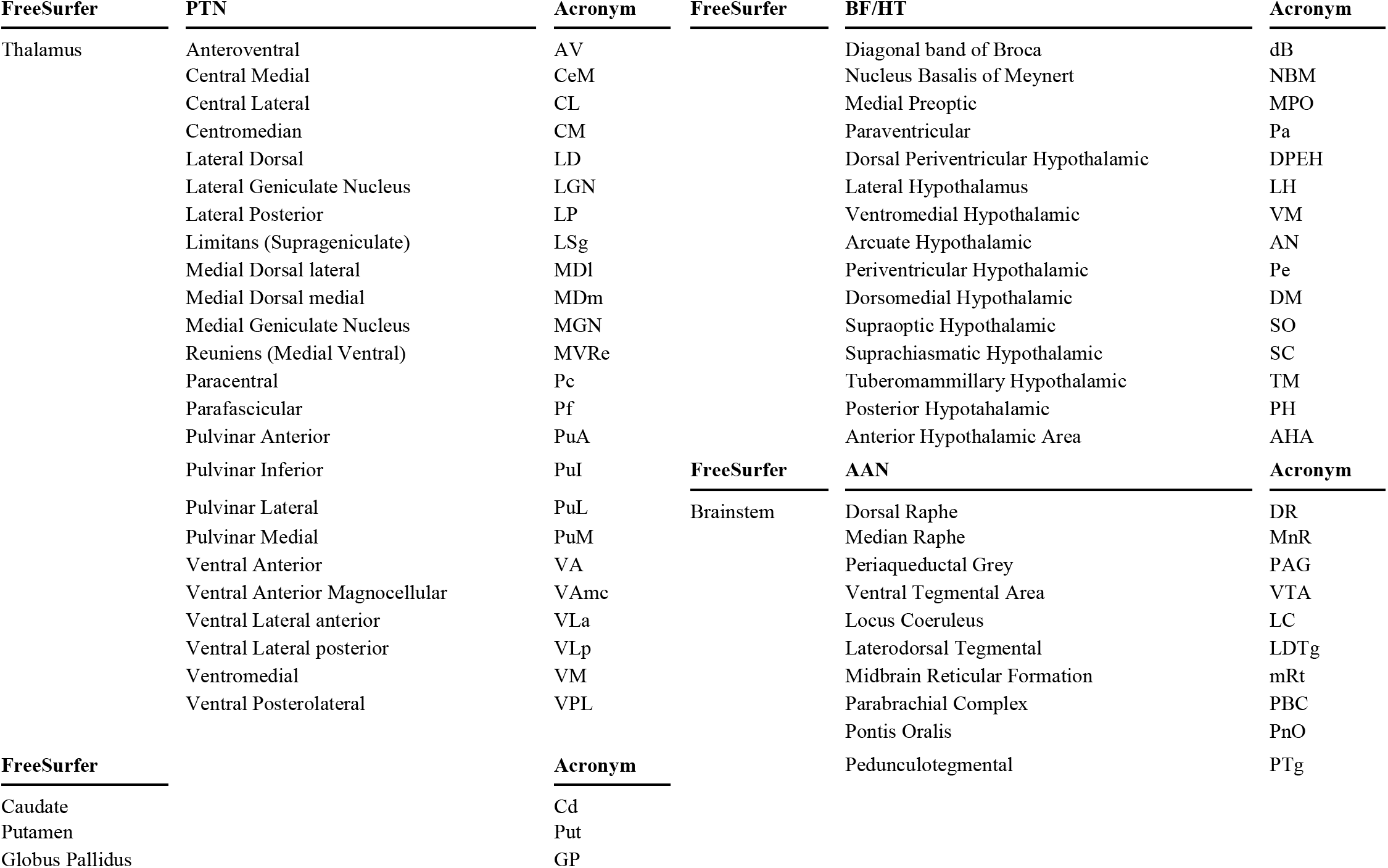
Subcortical regions of interest with respect to the FreeSurfer aseg altas (FreeSurfer), the probabilistic thalamic nuclei atlas (PTN), the Harvard ascending arousal network atlas (AAN), the basal forebrain / hypothalamus atlas (BF/HT), and their acronyms.

### 2.6. Quantitative analysis and hypothesis testing

Because the rs-fMRI time series were normalized to have zero mean and unit norm during preprocessing to satisfy the requirement for inter-subject synchronization (Joshi et al., 2018), the absolute values in the identified DMN are less interpretable than the relative differences among ROIs. Therefore, to facilitate a meaningful quantitative interpretation, we performed a normalization at each voxel of in the subcortical map by the 95% quantile (a scalar) of the values in the cortical DMN map, top of Fig. 1 (e). Thus, the normalized subcortical map indicates how strong the subcortical DMN activity is relative to the cortical DMN activity. Here we use the word “activity” to indicate the signals/values in either the cortical DMN map or subcortical DMN map (Fig. 1 (e)) identified using the NASCAR method. For each ROI, we plotted the normalized values of the subcortical map within that ROI, using a violin plot.

To address whether CL is strongly connected to the cortical DMN, we statistically tested if the mean of the subcortical DMN map within CL is significantly higher than that within the entire thalamus (including CL), using a two-sample t-test. Considering the volumetric interpolation procedure used above and the spatial smoothness of the rs-fMRI data, correction of *p*-values for multiple comparison are necessary. Corrections were performed based on random field theory (Brett et al., 2003; Worsley et al., 1992), where the number of resolution element (“resel”) was calculated based on the volume of each ROI and the FWHM of volumetric smoothing described in (Glasser et al., 2013). We similarly tested for VTA using this procedure, but with a comparison to the entire brainstem region (including VTA).

We then performed an exploratory study to test whether there are other candidate ROI(s) within the thalamus, hypothalamus, brainstem, and basal forebrain that have a strong connection to the cortical DMN, and thus could be used as subcortical targets for neuromodulation. We repeated the statistical testing above for all other ROIs defined in the PTN, AAN, and BF/HT atlas, with the proper correction for multiple comparisons as described above.

### 2.7. Comparison to seed-based method

We placed a single-vertex seed in the posterior cingulate cortex (PCC), which is one of the most commonly used seed locations in the DMN (Fox et al., 2005). For each voxel in the subcortical region, we then computed the Pearson correlation between that subcortical voxel and the seed, generating a seed-based subcortical functional connectivity map. Similar to the NASCAR analysis in Section 2.4, we visualized the result by overlaying it on the 100 micron structural dataset. We performed this seed-based analysis with and without global signal regression preprocessing. We repeated the above procedure using a second widely used single-vertex seed in the ventromedial prefrontal cortex (vmPFC).

### 2.8. Comparison of rs-fMRI results with brainstem immunostaining data

To validate the results, we compared the subcortical maps from the NASCAR and seed-based correlation analyses with tyrosine hydroxylase immunostain data from a human brainstem specimen. The brainstem specimen was donated from a 53-year-old woman, with written informed consent from a surrogate decision-maker as part of an Institutional Review Board-approved protocol. Additional details regarding the patient’s medical history, as well as the brainstem fixation and sectioning procedures, have been previously described, as this brainstem provided the basis for the Harvard AAN atlas used here (Edlow et al., 2012). For this analysis, we performed new tyrosine hydroxylase stains (rabbit polyclonal anti-tyrosine hydroxylase antibody; Pel-Freez Biologicals; Rogers AR) on tissue sections from the level of the caudal and rostral midbrain. Tyrosine hydroxylase stains dopamine-producing neurons, and thus was used as a reference standard for the accuracy of the VTA functional connectivity maps produced by the NASCAR and seed-based correlation analyses. The full tyrosine hydroxylase immunostaining protocol is available at https://github.com/ComaRecoveryLab/Subcortical_DMN_Functional_Connectivity and the stained tissue sections are available for interactive viewing at https://histopath.nmr.mgh.harvard.edu.

### 2.9. Reproducibility Analysis

We randomly split the 7T HCP rs-fMRI data into two halves and performed the same network identification procedure on these two independent datasets. The DMN was identified from each group and visually compared. Quantitatively, we also computed the Pearson correlation between the spatial map of the two identified DMNs.

## 3. Results

### 3.1. Qualitative visualization result

Fig. 2 (a) and (b) show the DMN subcortical map identified by the NASCAR method in an axial slice through the thalamus and striatum (for visualization only, NASCAR was running on the entire grayordinate data). The DMN subcortical map at the level of the basal forebrain, hypothalamus, and rostral midbrain is shown in Fig. 2 (c) and (d). Fig. 3 shows the DMN subcortical map at the level of the caudal midbrain in (a), (b), and the rostral pons in (c), (d). We use “correlation” hereafter to indicate the functional connectivity relationship of the subcortical regions to the cortical DMN. However, this “correlation” is not the Pearson correlation coefficient (see Section 3.3). Rather, it represents the strength of the DMN activity at each subcortical region. The higher the magnitude of the value in a subcortical region (the actual value could be either positive or negative), the stronger “resonance” of this region to the cortical DMN. The entire 3D volumetric results are shown as a video in the supplementary material and available at https://github.com/ComaRecoveryLab/Subcortical_DMN_Functional_Connectivity. The visualizat-ion results are not sufficient for making inferences at individual voxels due to the low spatial resolution of the fMRI data, as well as imperfect inter-subject coregistration (more detail in the Discussion section). Supplementary Fig. S1 includes the DMN functional connectivity map in its native resolution overlaid on the same 100 micron structural MRI for reference.

**Fig. 2.**
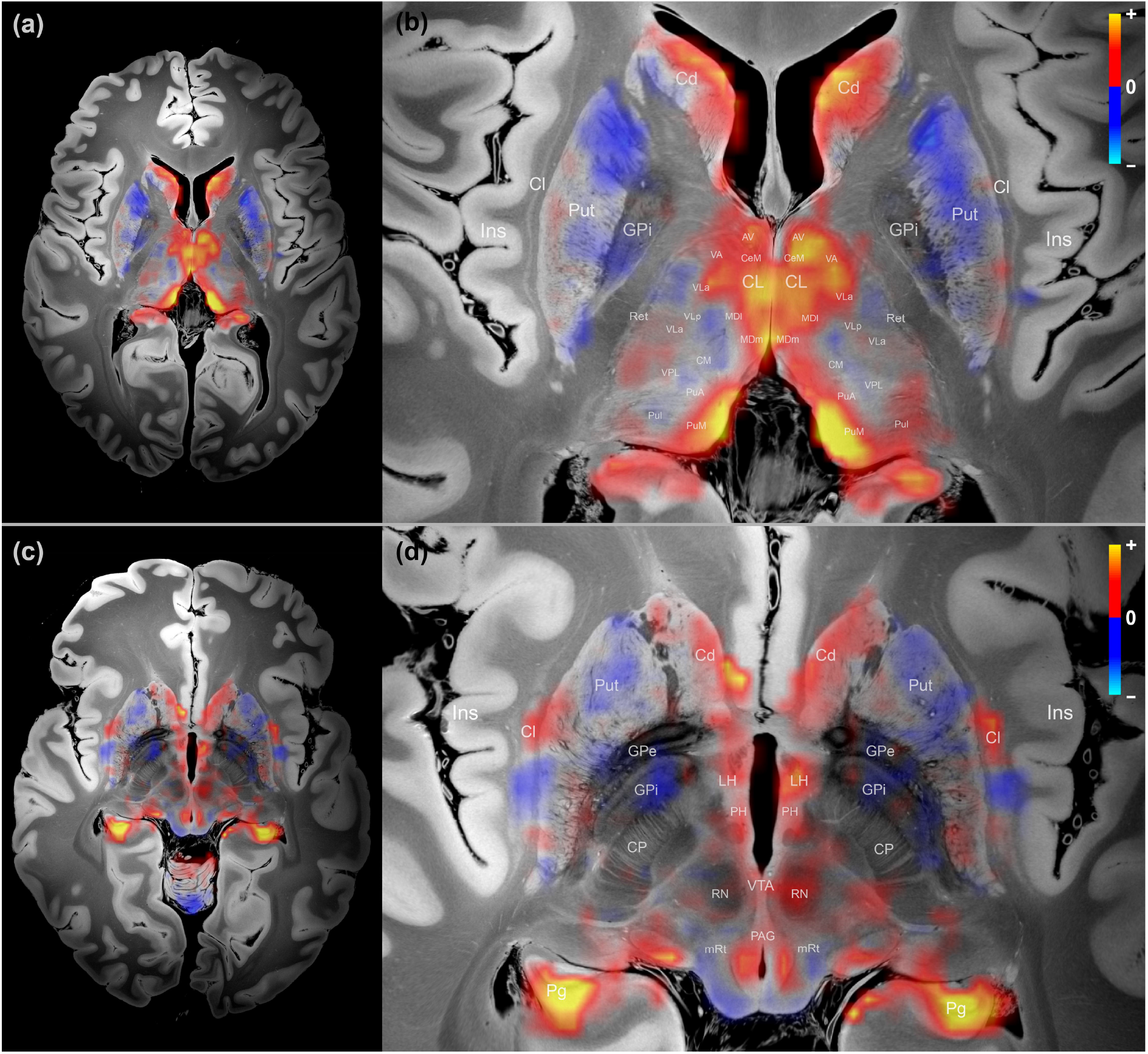
Map of the subcortical DMN connectome. (a) Overview of thalamus and basal ganglia; (b) Zoom-in version of (a) with annotations; (c) Overview of rostral midbrain; (d) Zoom-in version of (c) with annotations. The warm color (yellow/orange) indicates positive association or correlation with the DMN, and the cold color (blue) indicates negative association or anti-correlation with the DMN. Cl – Claustrum; CP – cerebral peduncle; GPe – Globus Pallidus Externus; GPi – Globus Pallidus Internus; Ins – Insula; Pg – parahippocampal gyrus; Ret – Reticular nuclei; Refer to Table 1 for other acronyms.

**Fig. 3.**
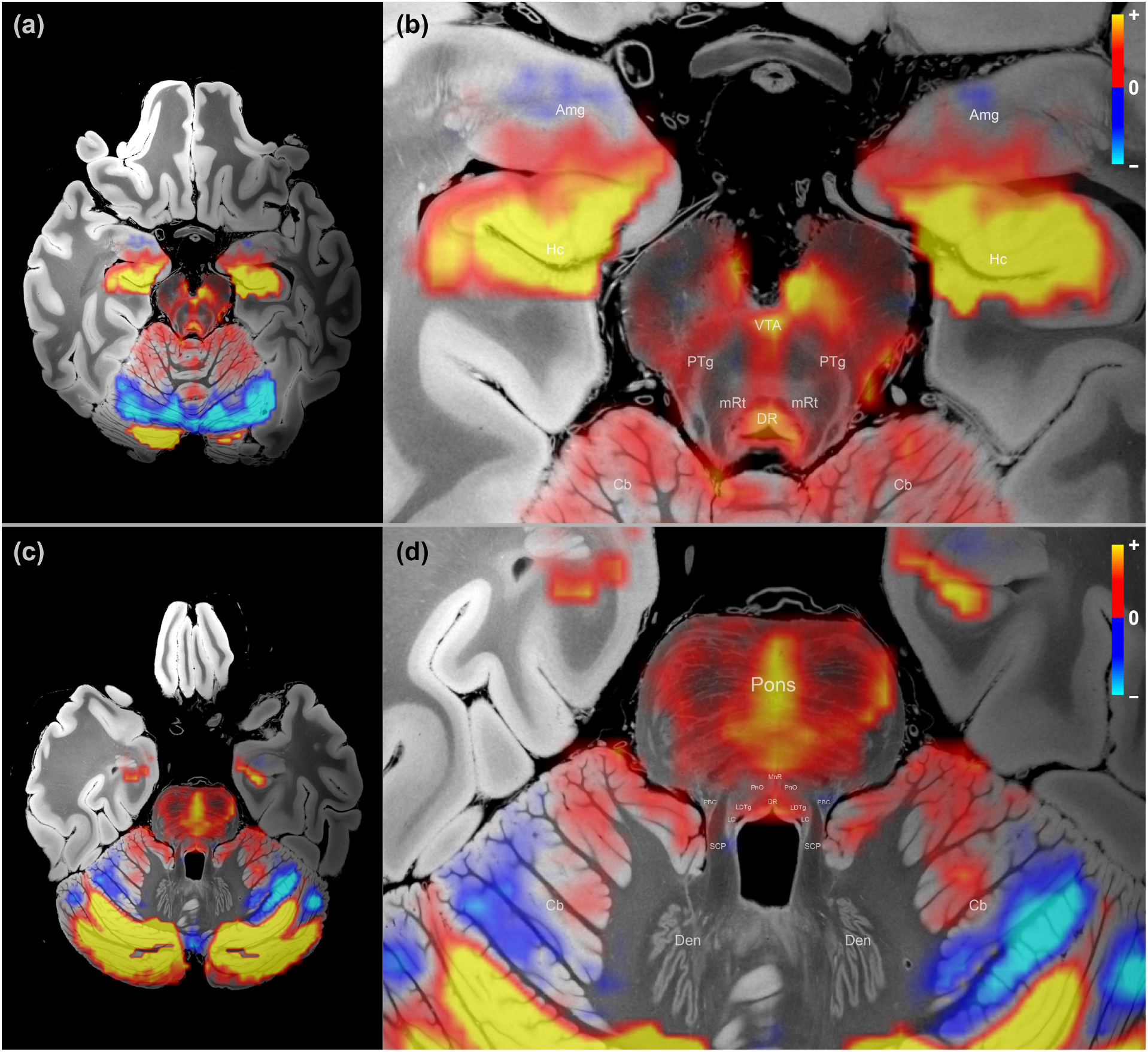
Map of the subcortical DMN connectome. (a) Overview of caudal midbrain; (b) Zoom-in version of (a) with annotations; (c) Overview of rostral pons; (d) Zoomed version of (c) with annotations. See Fig. 2 for color scheme. Cb – cerebellum; Den – dentate nucleus of the cerebellum; SCP – superior cerebellar peduncle. Refer to Table 1 for other acronyms.

Overall, we observed that the DMN subcortical components were largely symmetric about the midline of the brain, and the patterns appeared as spatially contiguous blobs. Importantly, there was no spatial constraint in the low-rank model itself, as shown in Fig. 1 (d), where each voxel/vertex was treated independently as part of the grayordinate representation during the decomposition. In other words, a random shuffle of the vertices/voxels before decomposition and a corresponding shuffle in the reverse order after the decomposition would not change the results, indicating that these resulting patterns likely reflect a physiological property of the data, not an artifact of the processing pipeline.

With respect to the neuroanatomic localization, the strongest regions of subcortical DMN functional connectivity were observed within the central thalamus, lateral hypothalamus, caudate nucleus, ventral tegmentum of the midbrain, periaqueductal grey area of the midbrain, and midline raphe of the midbrain and pons. All of these regions have animal or human evidence supporting their roles in the modulation of arousal, and hence consciousness (Alam et al., 2002; Eban-Rothschild et al., 2016; Lu et al., 2006; Parvizi, 2001; Van der Werf et al., 2002; Villablanca et al., 1976). The subcortical regions that showed the strongest anticorrelations with the DMN were the putamen and globus pallidus interna, regions that constitute the inhibitory component of a mesocircuit that was postulated to modulate the cerebral cortex via GABAerigc innervation of the central thalamus (Schiff, 2010). The basal forebrain did not contain large clusters of correlated or anti-correlated voxels.

### 3.2. Quantitative analysis result

Fig. 4 displays results of the analysis of subcortical functional connectivity with the DMN, using subcortical structures defined in the FreeSurfer aseg atlas. Subcortical regions are displayed along the x-axis, and the y-axis represents the normalized values with respect to the cortical DMN. We found that all subcortical structures exhibited substantially lower DMN activity compared to the cortex, with an averaged absolute percentage of 3.9%. We observed that the highest subcortical DMN signal is in the thalamus, caudate, and brainstem, reaching approximately 30% of the cortical DMN signal strength. The thalamus, caudate, brainstem, and hypothalamus showed strong positive correlations with the DMN, and the basal forebrain showed moderate positive correlations with the DMN, although negative correlations were also observed in these regions. Interestingly, the majority of voxels within the globus pallidus and putamen exhibited negative correlations with the DMN.

**Fig. 4.**
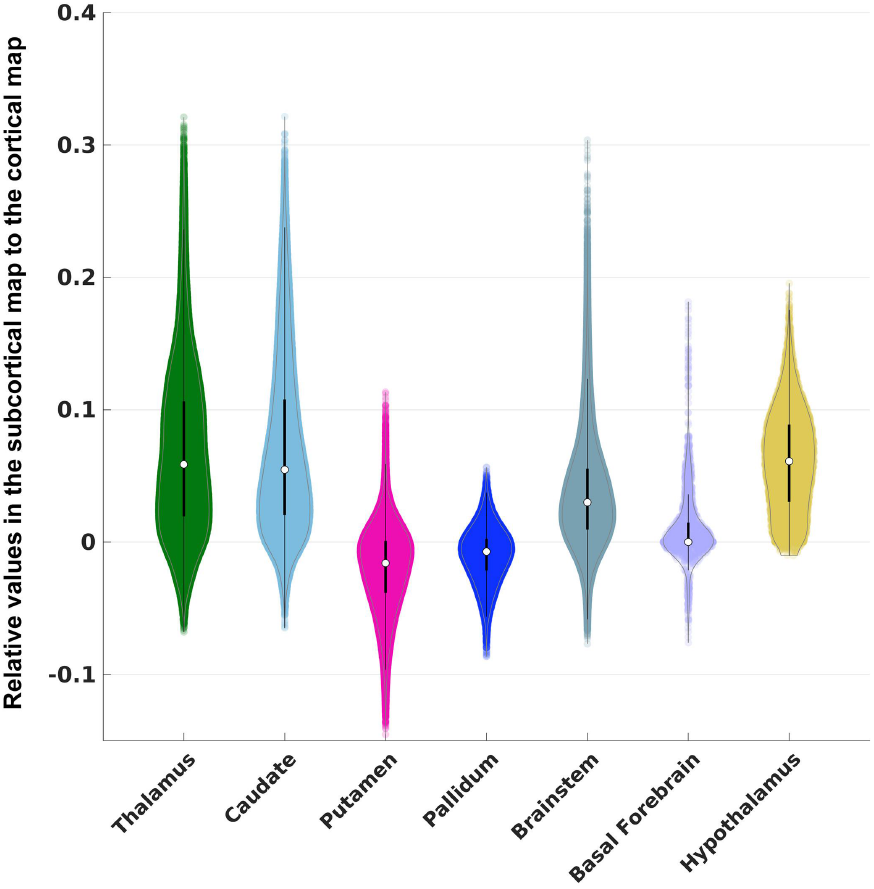
Violin plots for large-scale ROIs defined in FreeSurfer aseg atlas. Each violin plot shows the distribution of the DMN signals overlaid with dots for each individual voxel. The white dot indicates the median statistics and the black bar through the white dot is the traditional boxplot, where the thicker bar represents the 25% to 75% quantile and the thinner bar represents the whisker length that is 1.5 times of the interquartile, covering approximately 99.3% of the data range. The color scheme of the violin plots follows that in the atlas.

Fig. 5 shows the violin plots of the DMN signals for the thalamic nuclei. CL showed significantly higher functional connectivity with the cortical DMN than the average of thalamic signals. CL had the 4^th^ highest median value (slightly higher median values were observed in AV, MDm, and PuM), and the highest maximum value among all thalamic nuclei. CL contained voxels with the strongest correlation to the DMN, reflected by the heavy tail on top of its distribution.

**Fig. 5.**
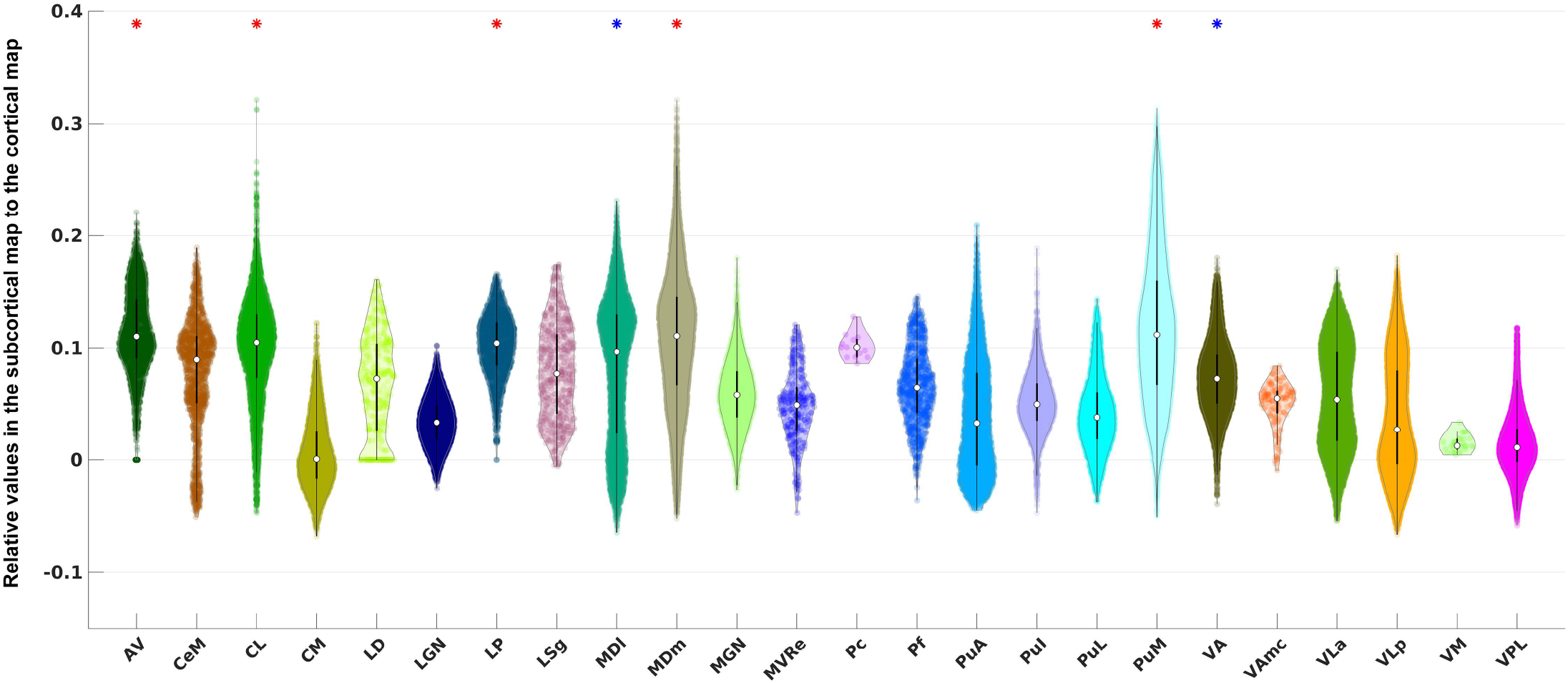
Violin plots for thalamic ROIs defined in the PTN atlas. The acronyms of the nuclei are labeled along the x-axis (see Table 1 for details). A star is placed above the violin plot if the average DMN signal within that nucleus is significantly higher than the mean signal of the entire thalamic region according to two sample Students’ t-test. The star is colored in blue if the *p*-value is below the standard *α* cutoff value of 0.05 after the correction and colored in red if the *p*-value is below an *α* cutoff value of 0.001 (this cutoff was chosen heuristically for contrasting and highlighting a higher significance).

Fig. 6 shows similar violin plots for the brainstem region. We found that the VTA had significantly higher DMN functional connectivity than the average of the brainstem, consistent with previous studies (Bär et al., 2016). Although VTA did not show the highest median value, it did contain the highest maximum value among all brainstem nuclei.

**Fig. 6.**
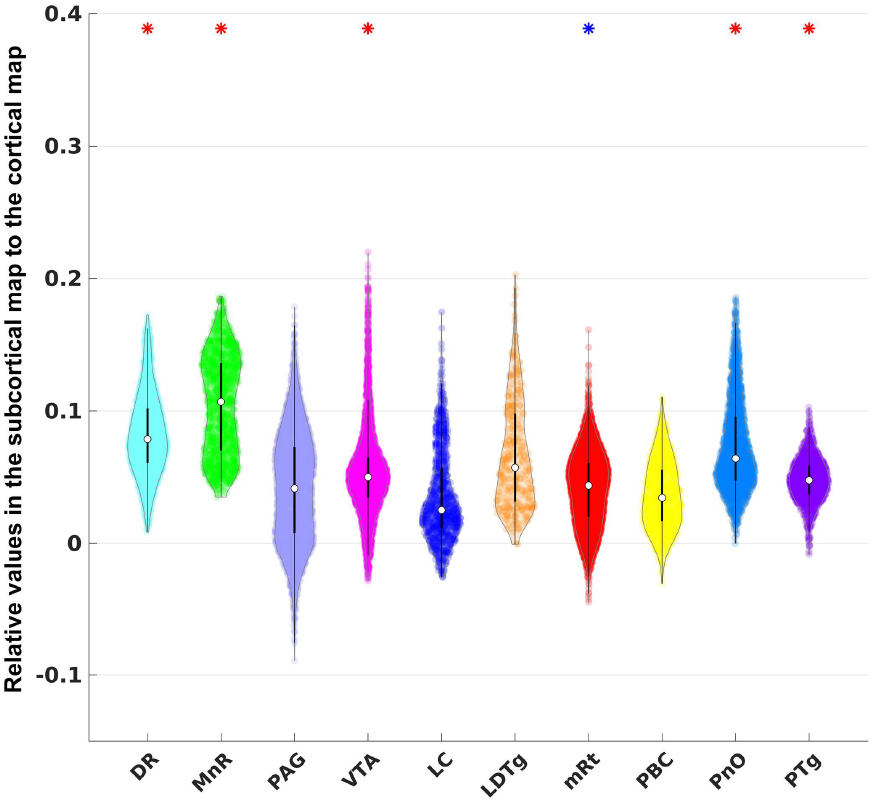
Violin plots for brainstem ROIs defined in the AAN atlas. See Fig. 5 for interpretation of the statistical significance.

In the exploratory analysis, we identified multiple additional nuclei that showed strong connection to the cortical DMN: AV, LP, MDl, MDm, PuM, and VA in the thalamus (Fig. 5), DR, MnR, mRt, PnO, and PTg in the brainstem (Fig. 6), and LH, VM, TM, and AHA in the hypothalamus (Fig. 7). Interestingly, distinct connectivity patterns were observed in dB and NBM, the two basal forebrain ROIs. Whereas dB showed exclusively positive DMN correlations, NBM showed a distribution of positive and negative correlations, yielding a median DMN connecting value close to zero. NBM was the only basal forebrain or hypothalamic nucleus with a substantial proportion of voxels showing negative correlations (i.e., anti-correlations) with the DMN.

**Fig. 7.**
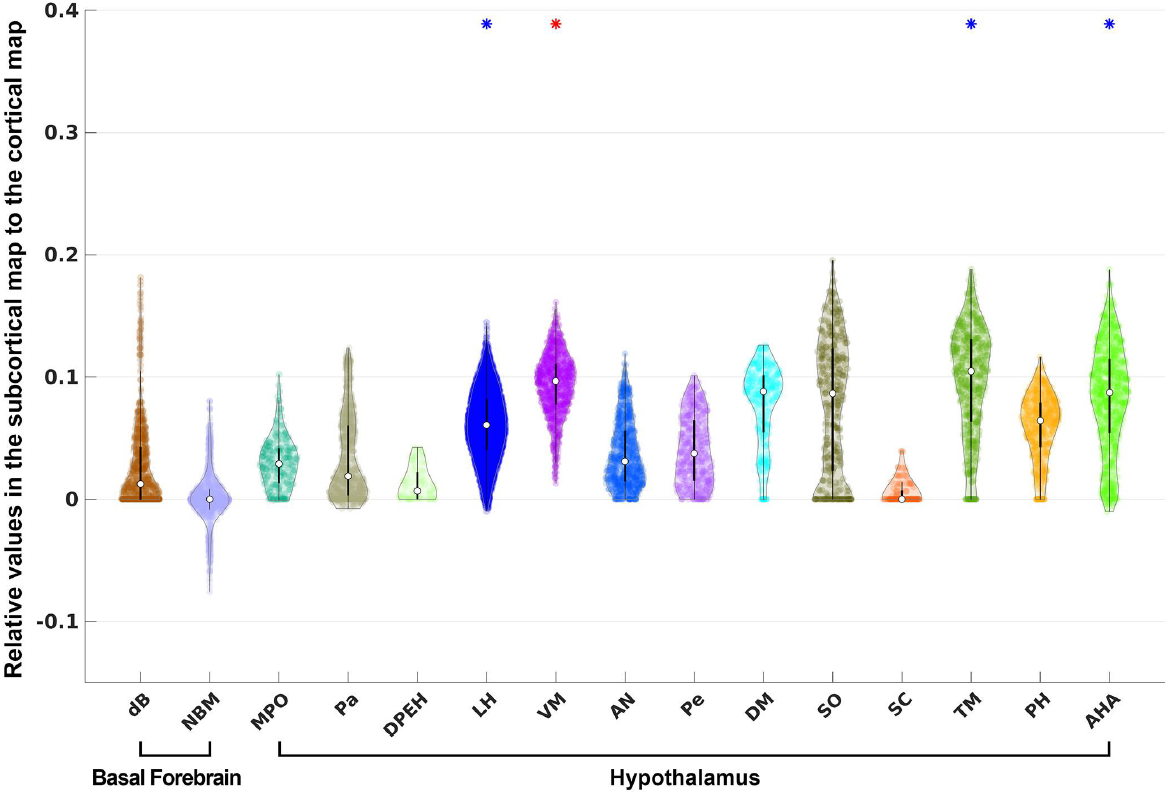
Violin plots for hypothalamus and basal forebrain ROIs defined in the atlas of Neudorfer *et al.,* 2020. See Fig. 5 for interpretation of the statistical significance.

### 3.3. Seed-based method and immunostaining images

Fig. 8 shows the seed-based correlation result for PCC in (a), (c) and for vmPFC in (b), (d) without global signal regression. The corresponding counterparts with global signal regression are shown in (e) – (h). Overall, these seed-based correlation results exhibit similar subcortical connectivity patterns (relative contrast between regions) to the patterns identified by NASCAR shown in Fig. 2 and Fig. 3. Specifically, in both the seed-based and NASCAR analyses, CL and VTA visually had the strongest correlation to the cortical DMN in comparison with other nuclei. However, the absolute correlation profile, especially the sign of the correlation (positively correlated vs anti-correlated), substantially differed between the results with global signal regression and in the ones without global signal regression. In the top row of Fig. 8, where no global signal regression had been applied, positive correlations were observed for most regions. This inflation of the correlation may be due to the global “physiological” or “vascular” component present in the fMRI data (Murphy and Fox, 2017; Zhu et al., 2015). In contrast, this “physiological” component was captured by NASCAR as a component separate from the DMN; thus, negative connections are clearly visible in the NASCAR result shown in Fig. 2 and Fig. 3. Although global signal regression can be used in seed-based analyses to reduce contamination from the “physiological” component, as shown in the bottom row of Fig. 8, it can be difficult to interpret the meaning of these negative correlations, as it has been shown mathematically that global signal regression introduces negative correlation into the seed-based correlation results (Murphy et al., 2009; Murphy and Fox, 2017). Indeed, the co-existence of both positive and negative correlation after the global signal regression in the dopaminergic VTA region, as shown in Fig. 8 (g) and (h), does not appear to be anatomically consistent with prior anatomic atlases (Edlow et al., 2012; Trutti et al., 2021), with our immunostain data (Fig. 9), or with prior neuronal labeling studies in rodents, non-human primates, and humans (Breton et al., 2019; Root et al., 2016; Taylor et al., 2014), as discussed below. Finally, there were substantial variations in the seed-based correlation result depending on the choice of the seed point, which is consistent with findings in the literature (Uddin et al., 2009). For example, negative correlations were observed in the putamen and globus pallidus when PCC was selected as the seed shown in Fig. 8 (a), whereas they were barely visible when vmPFC was used as the seed shown in Fig. 8 (b).

**Fig. 8.**
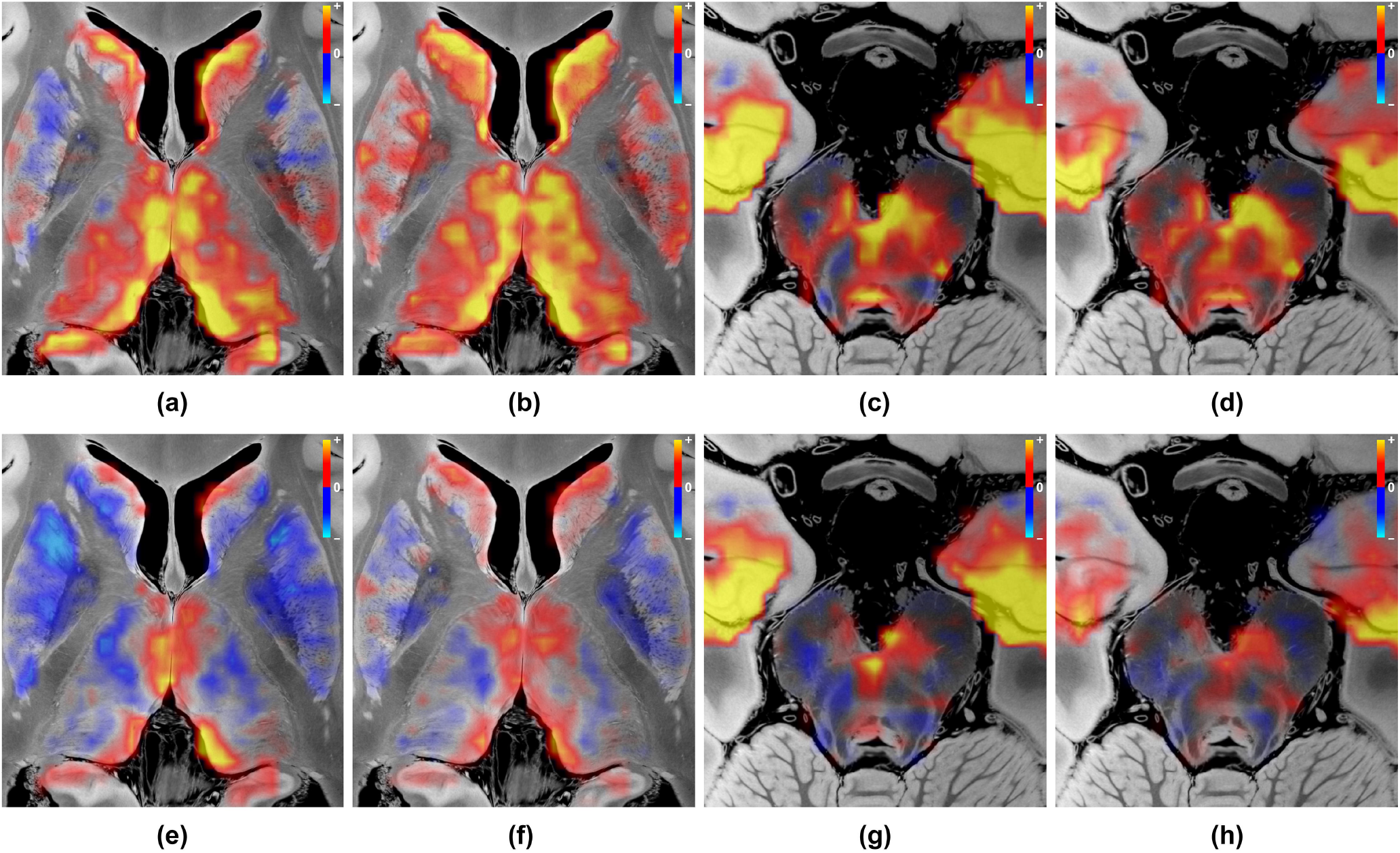
Seed-based correlation analysis results. (a) PCC-seeded correlation map in the same axial plane as Fig. 2 (b), showing correlation structures in thalamus and basal ganglia without global signal regression; (b) Same as (a) but using the seed point in vmPFC; (c) Same as (a) but in the caudal midbrain plane, Fig. 3 (b); (d) Same as (c) but using the seed point in vmPFC; (e) – (h) Same as (a) – (d) but with global signal regression.

**Fig. 9.**
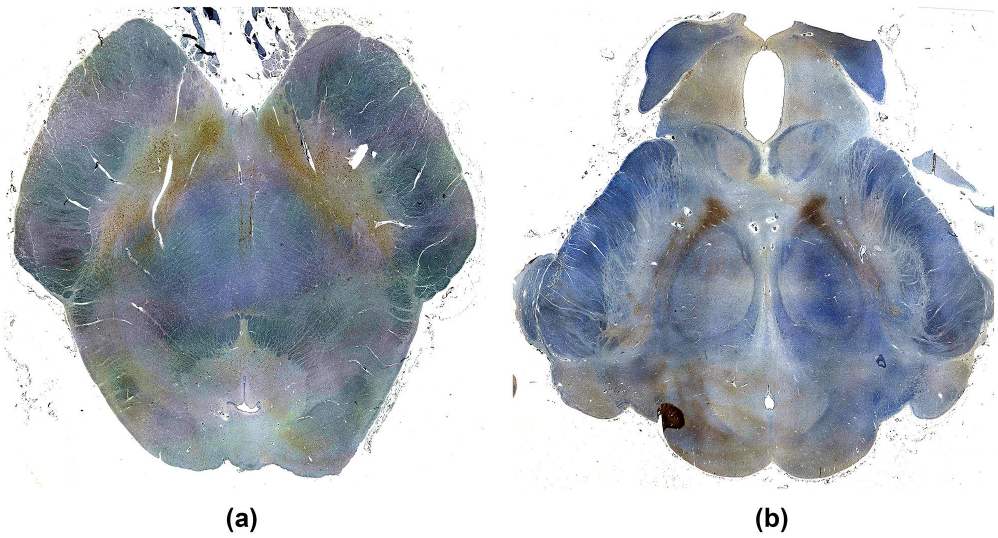
Immunostain images of dopaminergic ventral tegmental area (VTA) neurons in (a) the caudal midbrain and (b) the rostral midbrain. Dopaminergic neurons (brown) were immunostained with tyrosine hydroxylase, and each axial section was counterstained with hematoxylin (blue). In both the caudal and rostral midbrain sections, the VTA neurons extend laterally and posteriorly along the lateral border of the decussation of the superior cerebellar peduncles (a) and the red nuclei (b). The human brainstem specimen used for these immunostains was donated by a 53-year-old woman who died of non-neurological causes. A surrogate-decision maker provided written informed consent for brain donation and postmortem research. Additional details about the specimen have been previously published (Edlow et al. 2012).

### 3.4. Reproducibility Analysis

The spatial maps of the identified DMN from the two independent groups were visually indistinguishable, and the correlation coefficient between the two spatial maps (grayordinate representation) was 0.987, demonstrating the high reproducibility of the results using our tensor-based analysis pipeline.

## 4. Discussion

In this brain mapping study using 7T resting-state fMRI data from the HCP, we characterized the subcortical connectivity of the human DMN and openly release the results in standardized form (via <u>Lead-DBS</u>, <u>Openneuro</u>, and <u>FreeSurfer</u>). We expanded the DMN to include subcortical nodes by a joint analysis on the grayordinate system, providing a more complete map of the human DMN. The map was generated by a tensor-based NASCAR decomposition method that revealed extensive interconnections between the canonical cortical DMN and subcortical regions within the brainstem, thalamus, hypothalamus, basal forebrain, and basal ganglia. Further, the NASCAR and seed-based correlation results supported our hypothesis that CL and VTA are subcortical nodes strongly connected to the cortical DMN. The spatial, temporal, and physiological properties (e.g., correlations versus anti-correlations) of the subcortical DMN connectivity map create new opportunities to elucidate subcortical contributions to human consciousness and provide therapeutic targets for interventions that aim to promote recovery of consciousness in patients with severe brain injuries.

Our functional connectivity results are consistent with, and build upon, decades of electrophysiological and neuroimaging investigations of the subcortical networks that modulate consciousness. For example, the CL nucleus of the thalamus is a well-established hub node of reciprocal thalamocortical networks, as CL is richly innervated by arousal neurons of the brainstem and basal forebrain (Steriade and Glenn, 1982) and provides diffuse innervation of the neocortex (Dempsey and Morison, 1941). More recently, deep brain stimulation studies targeting CL in non-human primates (Baker et al., 2016; Redinbaugh et al., 2020) and humans with severe brain injuries (Schiff et al., 2007) have confirmed the role of CL in modulating consciousness. However, a non-invasive, fMRI-based biomarker of CL functional connectivity has been elusive, and the mechanisms and pathways by which the central thalamus modulates the DMN remain an area of active study (Cunningham et al., 2017; Guldenmund et al., 2013; Wang et al., 2014). Here, we provide robust evidence, with out-of-sample testing, for strong positive correlations between the CL nucleus and the human DMN. Indeed, individual voxels within CL showed the strongest correlation with the DMN of any thalamic voxels, and the median correlation strength of CL voxels with the DMN was the 4^th^ highest of all thalamic nuclei. These observations suggest that high spatial and temporal resolution rs-fMRI with advanced signal processing and modeling methods, as performed here, provide a potential biomarker of CL-DMN functional connectivity and an opportunity to test the hypothesis that CL stimulation induces reemergence of consciousness via CL-DMN functional connectivity in patients with severe brain injuries.

The potential translational impact of the CL-DMN functional connectivity findings is particularly noteworthy when considered in the context of the mesocircuit hypothesis of consciousness, for which we provide new rs-fMRI functional connectivity evidence in the human brain. Specifically, we observed anti-correlations between the putamen, globus pallidus, and the DMN, indicating that putaminal and pallidal activity toggles inversely with the thalamic CL and the cortical DMN. These observations are consistent with known GABAergic inhibitory neuronal inputs from GPi to CL, a neuroanatomic relationship that has been suggested by clinical case studies but has not been directly observed with fMRI in the human brain. Since the proposal of the mesocircuit hypothesis in 2010 (Schiff, 2010), multiple clinical observations have supported the hypothesis (Fridman et al., 2014; Williams et al., 2013), including the paradoxical therapeutic response of approximately 5% of patients with severe brain injuries to the GABAergic medication zolpidem (Whyte et al., 2014) – a response believed to be mediated by restoration of CL disinhibition within the mesocircuit. Our subcortical functional DMN connectivity results raise the possibility that individualized rs-fMRI maps of GPi-CL-DMN functional connectivity can be used in future clinical trials as a predictive biomarker (i.e., to identify patients who are likely responders to GABAergic therapy) and as a pharmacodynamic biomarker (i.e., to test whether a therapy engages the mesocircuit target).

Importantly, BOLD fMRI anti-correlations recorded at mesoscale should not be interpreted as a direct measure of neuronal signals recorded at microscale (Fox et al., 2009). Anti-correlations reflect numerous neurovascular-glial properties (Fox et al., 2009; He et al., 2018), and thus our anti-correlation results do not provide direct proof of the mesocircuit hypothesis. Nevertheless, our rs-fMRI methods provide a mesoscale biomarker of mesocircuit integrity that may have utility in clinical trials, particularly when considering that non-invasive stimulation techniques such as LIFUP are now targeting the globus pallidus (Cain et al., 2021b) and central thalamus (Cain et al., 2021a; Monti et al., 2016) in patients with disorders of consciousness.

Additional neuroanatomic insights provided by the subcortical DMN connectivity map include new evidence for brainstem nodes that are strongly connected to the DMN. The VTA functional connectivity findings confirmed our hypothesis that the VTA is strongly connected to the DMN, consistent with recent rs-fMRI evidence for VTA functional connectivity with the posterior cingulate/precuneus, a central hub node of the DMN (Buckner and DiNicola, 2019; Thomas Yeo et al., 2011). VTA modulation of consciousness via dopaminergic signaling has been suggested by preclinical studies using pharmacologic (Kenny et al., 2015; Solt et al., 2011), electrical (Solt et al., 2014), optogenetic (Eban-Rothschild et al., 2016; Taylor et al., 2016), and chemogenetic (Oishi et al., 2017) stimulation, as well as a mouse dopamine knock-out model (Palmiter, 2011). However, until recently, there has only been indirect evidence for dopaminergic VTA modulation of human consciousness from pharmacological studies using dopaminergic drugs (Fridman et al., 2019, 2010; Giacino et al., 2012), as well as positron emission tomography studies of dopamine receptor dynamics (Fridman et al.,2019). Now, with emerging evidence for dopaminergic VTA modulation of human consciousness via DMN functional connectivity, there is a compelling clinical need for robust and reliable biomarkers of VTA-DMN functional connectivity. A major goal for future research will be to determine if such a biomarker can be validated on 3T scans that are used for clinical purposes.

Beyond CL and VTA, our exploratory analyses suggest that the DMN has subcortical connections in additional regions of the brainstem, hypothalamus, thalamus, basal ganglia, and basal forebrain. These findings should be considered hypothesis-generating and will require validation in future connectivity studies. To inform the design of future experiments, we emphasize that several subcortical nuclei that demonstrated DMN correlations have strong data to support their role in modulating arousal, and hence consciousness, in prior animal studies. In particular, DR and MnR have been shown in animal electrophysiological experiments to regulate arousal (McGinty and Harper, 1976; Trulson and Jacobs, 1979; Xi et al., 2004), and these serotonergic nuclei in the pontomesencephalic raphe have been shown to be functionally connected to the DMN (Bär et al., 2016) or to constitute a sub-network of the DMN (Beliveau et al., 2015) in prior human rs-fMRI experiments. The mRt and PnO nuclei, classically considered the brainstem’s “reticular core”, are also recognized as key nodes of an ascending reticular activating system, based on decades of electrophysiologic investigations of arousal in animal models (Moruzzi and Magoun, 1949; Steriade et al., 1982; Xi et al., 2004). Similarly, our observation of lateral hypothalamic area and tuberomammillary nucleus connectivity with the DMN is consistent with electrophysiologic studies in rodent models of sleepwake cycle regulation (Alam et al., 2002; Takahashi et al., 2006). Collectively, these exploratory results thus add to a strong body of evidence in animal models, and a small but growing body of evidence in human studies, that multiple subcortical regions are involved in arousal regulation. Our findings expand upon prior studies by suggesting that DMN functional connectivity is a mechanism by which these subcortical nuclei activate the cerebral cortex to promote consciousness. Elucidation of the precise physiological mechanisms, temporal dynamics, and anatomic subspecialization of these subcortico-cortical connections will require multi-modality investigations of arousal in animal models (Pais-Roldán et al., 2019) and human experiments (Fultz et al., 2019) designed to interrogate the structure and function of subcortical arousal pathways at increasingly high levels of spatial and temporal resolution (DeFelipe, 2010).

When averaged over data from many subjects, the NASCAR and seed-based correlation methods yielded similar spatial patterns and contrast of the subcortical DMN. There has been debate about whether global signal regression helps or hurts the correlation analysis (Murphy and Fox, 2017). Without global signal regression, the correlation measures tend to be inflated due to the involvement of global “physiological” signals (Chen et al.,2020). On the other hand, the application of global signal regression introduces negative correlations (Murphy et al., 2009; Murphy and Fox, 2017). In contrast, NASCAR directly models the “global physiological” network as one of the low-rank components. In fact, this component was identified as the first network during the tensor decomposition, with the network strength *λ* even higher than that of the DMN. In this way, NASCAR successfully decoupled the DMN from this global component, avoiding the ambiguity in the interpretation of the seed-based correlation results.

A manifestation of this ambiguity generated by seed-based correlation with global signal regression is seen in the VTA functional connectivity results. Whereas the NASCAR method detected DMN correlations within the VTA, the seed-based correlation with global signal regression detected correlations and anti-correlations. Prior studies have shown that the VTA contains not only dopaminergic neurons, but also GABAergic and glutamatergic neurons, raising the possibility that VTA interactions with the DMN could be spatially heterogeneous. However, prior neuronal labeling studies in rodents, non-human primates, and humans (Breton et al., 2019; Root et al., 2016; Taylor et al., 2014) indicate that dopaminergic, GABAergic, and glutamatergic neurons are intermingled within the VTA, making it unlikely that there would be discrete subregions of BOLD correlations and anticorrelations within the VTA. Furthermore, our immunostaining results (Fig. 9) revealed a symmetric, spatially contiguous distribution of dopaminergic neurons within the VTA. Given that the majority of VTA neurons produce dopamine (Taylor et al., 2014), these immunostaining results suggest that the spatially contiguous correlation results generated by the NASCAR method are more anatomically plausible than the spatially disparate correlation and anti-correlation results generated by the seed-based correlation with global signal regression.

A limitation of the NASCAR approach is the assumption of perfect inter-subject coregistration. The low-rank tensor model would fail if there were substantial spatial misalignment among subjects. In this study, we relied on the boundary-based registration method (Greve and Fischl, 2009) used in the HCP minimal preprocessing pipeline (Glasser et al., 2013). Although many model-based and more recently deep-learning-based approaches (Balakrishnan et al., 2019; Cheng et al., 2020; Robinson et al., 2014; Yeo et al., 2010) have been proposed to improve inter-subject coregistration, registration of subcortical regions remains challenging, especially when working with fMRI data where the spatial resolution is insufficient. For these reasons, although our results shown in Figs. 2 and 3 exhibited neuroanatomically plausible patterns of the subcortical DMN, we caution against making inferences on the voxel level, particularly near the boundaries of subcortical structures. The functional connectivity map in supplementary Fig. S1 shows the functional connectivity results before interpolation and illustrates this limitation.

As with other data-driven approaches (e.g., ICA), post-hoc manual inspection of the decomposed components is required. Theoretically, one NASCAR component may contain multiple networks, or multiple NASCAR components may represent sub-networks of a large-scale canonical network. However, in this work, the DMN was identified as the second component and confirmed by visualization of the cortical map spatially shown in Fig. 1 (e). We did not find other components exhibiting a spatial map similar to the DMN. Due to this data-driven property of the NASCAR decomposition method, exploration of other components besides the DMN is a promising future direction.

The SNR of fMRI data is very low, as all 30 networks extracted using the NASCAR method accumulatively could only explain ~11% of the variance in the data. Moreover, the SNR in subcortical regions is much lower than that in the cerebral cortex, making it difficult to identify functional connectivity in subcortical regions, particularly in the brainstem (Sclocco et al., 2018). As seen in Fig. 4, the average magnitude of DMN signals identified in the subcortical regions is only ~5% of that in the cortical nodes. We attempted to address this inherent limitation of subcortical fMRI data by analyzing the 7T HCP rs-fMRI dataset, which has possibly the best SNR of any publicly available dataset. Hence, it is difficult to judge whether the results discovered in this work will be reproducible when the method is applied to datasets with lower SNR, e.g., HCP 3T or clinical 3T scans. Reproducing and generalizing these findings to 3T scanners and lower resolution rs-fMRI datasets is essential for clinical translation and is a key goal for future investigation.

## 5. Conclusion

We provide a functional connectivity map of the subcortical DMN in the human brain. We reveal new functional connectivity properties of the brainstem, hypothalamus, thalamus, and basal forebrain, which may be used in future investigations of subcortical contributions to human consciousness. The subcortical DMN connectivity map may also be used in clinical trials as a predictive biomarker to inform patient selection or as a pharmacodynamic biomarker to measure whether a therapy engages its target within the DMN. We release the subcortical DMN connectivity map via <u>Lead-DBS</u>, <u>FreeSurfer</u> and <u>Openneuro</u> platforms for use in future neuromodulation studies.

## Data and code availability

The data used in this study are publicly available from the Wash U/U Minn component of the Human Connectome Project, Young Adult Study at https://www.humanconnectome.org/study/hcp-young-adult.

For research purposes, we release the subcortical functional map of the default mode network at Lead-DBS (https://www.lead-dbs.org/helpsupport/knowledge-base/atlasesresources/atlases), FreeSurfer (https://surfer.nmr.mgh.harvard.edu), and OpenNEURO (https://openneuro.org/datasets/ds003716), and the open-access code is released at the GitHub repository (https://github.com/ComaRecoveryLab/Subcortical_DMN_Functional_Connectivity).

## Acknowledgment

This study was supported by the NIH Director’s Office (DP2HD101400), National Institute of Neurological Disorders and Stroke (R21NS109627, RF1NS115268, R01NS0525851, R21NS072652, R01NS070963, R01NS083534, U01NS086625, U24NS10059103, R01NS105820), BRAIN Initiative Cell Census Network (U01MH117023), National Institute for Biomedical Imaging and Bioengineering (P41EB015896, R01EB023281, R01EB006758, R21EB018907, R01EB019956, P41EB030006), National Institute on Aging (R56AG064027, R01AG064027, R01AG008122, R01AG016495), National Institute of Mental Health (R01MH123195, R01MH121885, 1RF1MH123195), the James S. McDonnell Foundation, and the Tiny Blue Dot Foundation. This work was also made possible by the resources provided by Shared Instrumentation Grants S10RR023401, S10RR019307, and S10RR023043. Additional support was provided by the NIH Blueprint for Neuroscience Research (U01MH093765), part of the multi-institutional Human Connectome Project.

