## Supplementary Material for "Mapping the subcortical connectivity of the human default mode network"

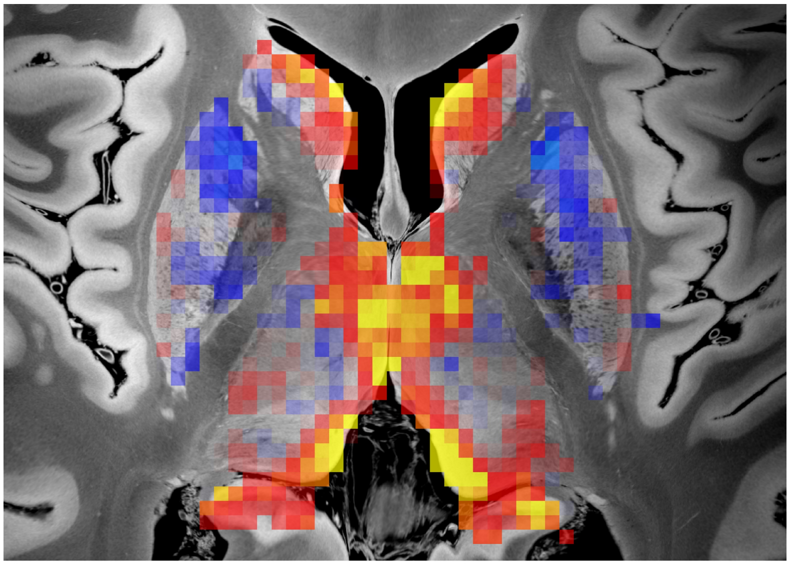

**(a)**

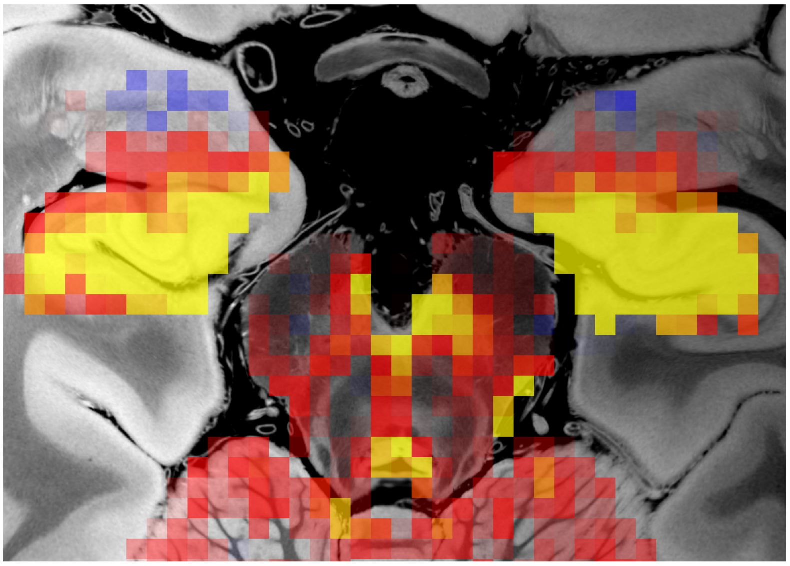

**(b)**

**Fig. S1.** Subcortical map of the default mode network obtained using the Nadam-accelerated SCAlable and Robust tensor decomposition method in the original resting-state functional MRI resolution. Representative images are shown in the axial plane at the level of the (a) mid-thalamus and (b) caudal midbrain.
